# Taking the next-gen step: comprehensive antimicrobial resistance detection from *Burkholderia pseudomallei* genomes

**DOI:** 10.1101/720607

**Authors:** Danielle E. Madden, Jessica R. Webb, Eike J. Steinig, Bart J. Currie, Erin P. Price, Derek S. Sarovich

## Abstract

**Background:** Antimicrobial resistance (AMR) poses a major threat to human health. Whole-genome sequencing holds great potential for AMR identification; however, there remain major gaps in comprehensively detecting AMR across the spectrum of AMR-conferring determinants and pathogens.

**Methods:** Using 16 wild-type *Burkholderia pseudomallei* and 25 with acquired AMR, we first assessed the performance of existing AMR software (ARIBA and CARD) for detecting clinically relevant AMR in this pathogen. *B. pseudomallei* was chosen due to limited treatment options, high fatality rate, and AMR caused exclusively by chromosomal mutation (i.e. single-nucleotide polymorphisms [SNPs], insertions-deletions [indels], copy-number variations [CNVs], and functional gene loss). Due to poor performance with existing tools, we developed ARDaP (<u>A</u>ntimicrobial <u>R</u>esistance <u>D</u>etection <u>a</u>nd <u>P</u>rediction) to identify the spectrum of AMR-conferring determinants in *B. pseudomallei*.

**Results:** CARD failed to identify any clinically-relevant AMR in *B. pseudomallei*, ARIBA cannot differentiate AMR determinants from natural genetic variation, and neither CARD or ARIBA can identify CNV or gene loss determinants. In contrast, ARDaP accurately detected all SNP, indel, CNV, and gene loss AMR determinants described in *B. pseudomallei* (*n*≈50). Additionally, ARDaP accurately predicted three previously undescribed determinants. In mixed strain data, ARDaP identified AMR to as low as ~5% allelic frequency.

**Conclusions:** We demonstrate that existing AMR software are inadequate for comprehensive AMR detection; ARDaP overcomes the shortcomings of existing tools. Further, ARDaP enables AMR prediction from mixed sequence data down to 5% allelic frequency. ARDaP databases can be constructed for any microbial species of interest for comprehensive AMR detection.

## Introduction

Antimicrobial resistance (AMR) poses a major threat to human health worldwide and an increasing contributor to morbidity and mortality. Antibiotic use and misuse have resulted in an alarming increase in multidrug-resistant infections worldwide, resulting in an urgent need to improve global AMR detection and surveillance. Alongside pathogen identification, AMR detection is one of the primary goals of diagnostic microbiology, with far-reaching consequences for both infection control and effective treatment [1].

Whole-genome sequencing (WGS) holds great potential for comprehensive AMR detection and prediction from bacterial genomes by identifying all AMR determinants in a single genome or metagenome [2], circumventing the need for multiple and often laborious diagnostic methods. Existing bioinformatic tools such as ARG-ANNOT [3], Antibiotic Resistance Identification By Assembly (ARIBA) [4], Comprehensive Antibiotic Resistance Database (CARD) [5], and MEGARes [2] can readily detect AMR genes acquired from horizontal gene transfer events. Many bacterial pathogens also develop AMR via chromosomal mutations, including missense single-nucleotide polymorphism (SNP) mutations in β-lactamase-encoding genes, SNPs or nonsense insertion-deletions (indels) in efflux pump regulators [6–8], gene amplification via copy-number variations (CNVs) [9], and functional gene loss [6]. Recent improvements in AMR identification software mean that chromosomal mutations, particularly SNPs, are now identifiable. For example, ARIBA can identify AMR-conferring SNPs and indels in multiple species [4]. Nevertheless, other types of genetic variants −particularly gene loss or truncation, inversions, and CNVs – remain poorly identified using existing tools, despite their crucial role in conferring AMR [10].

The bacterium, *Burkholderia pseudomallei*, causes the often-fatal tropical disease melioidosis. Melioidosis severity ranges from mild, self-limiting skin abscesses to pneumonia, neurological disease, and septic shock. *B. pseudomallei* is naturally resistant to many antibiotics, including aminoglycosides, penicillins, macrolides, and polymyxins [11, 12]. Fortunately, human-to-human *B. pseudomallei* transmission is rare; almost all infections are acquired from the environment. As such, isolates collected prior to antibiotic treatment are almost universally susceptible to the following clinically-relevant antibiotics: ceftazidime (CAZ), amoxicillin-clavulanate (AMC), co-trimoxazole (SXT), doxycycline (DOX), meropenem (MEM) and imipenem (IPM) [13]. To prevent melioidosis relapse, treatment involves prolonged (3-6 month) antibiotic therapy, which increases AMR risk and treatment failure [6]. AMR in *B. pseudomallei* has been reported for all clinically-relevant antibiotics [6], with novel AMR determinants towards these key antibiotics continuing to be uncovered.

Here, we tested 47 characterised *B. pseudomallei* genomes with known antibiotic phenotype profiles and associated AMR determinants, and three MEM-resistant (MEMr) strains with previously unidentified AMR determinants, against existing tools (ARIBA and CARD) to determine their AMR detection efficacy. Among the characterised strains, 25 were phenotypically-confirmed as resistant towards at least one clinically relevant antibiotic, 16 were sensitive, and the remainder encoded unusual sensitivity towards aminoglycosides and macrolides or stepwise AMR variants. Following testing against the current AMR tools, we developed a new tool, <u>A</u>ntibiotic <u>R</u>esistance <u>D</u>etection <u>a</u>nd <u>P</u>rediction (ARDaP), to permit comprehensive AMR detection from microbial genomes. ARDaP was designed to meet four main aims: first, to accurately identify AMR determinants caused by a spectrum of mutational mechanisms (i.e. gene gain, SNPs, indels, CNVs, and functional gene loss); second, to predict enigmatic AMR determinants in isolates with phenotypically-confirmed AMR, third, to detect minor AMR allelic determinants in mixed (e.g. metagenomic) sequence data; and finally, to provide a user-friendly report that summarises the AMR determinants (if any) and associated AMR phenotypes, stepwise variants, unusual antimicrobial sensitivity determinants, and genetic variants associated with natural variation that do not confer AMR. Although we illustrate its utility in *B. pseudomallei*, ARDaP is amenable to AMR identification across all microbial species.

## Methods

### Ethics

Ethical approval was obtained as previously described [14].

### Isolates

Forty-seven *B. pseudomallei* strains were included in this study, including 25 with elevated MICs towards one or more clinically-relevant antibiotics (Table 1) and genotypically-confirmed AMR determinants. These isolates were selected as they represent the spectrum of known AMR determinants in *B. pseudomallei* (Table S1). Strains encoding unusual aminoglycoside- and macrolide-sensitivity, and stepwise mutations that lower the barrier to AMR development, were also examined (Table 1). A further 16 strains sensitive to all clinically-relevant antibiotics were included to test software efficacy (Table 2). Finally, three previously uncharacterised clinical strains exhibiting MEMr (MSHR1058 MIC=12 μg/mL; MSHR1174 MIC=6 μg/mL; MSHR8777 MIC=4 μg/mL; Table 3) were included to test the predictive capacity of ARDaP.

**Table 1.** Antimicrobial resistance (AMR) determinants in 25 *Burkholderia pseudomallei* strains with verified AMR phenotypes, plus strains conferring unusual antimicrobial susceptibility and stepwise AMR variants^§^

| Patient ID | Isolate | ST | Genome accession | Antibiotic MIC (µg/mL)<sup>‡</sup> | Stepwise variant<sup>†</sup> | AMR determinant | Reference/s |
| --- | --- | --- | --- | --- | --- | --- | --- |
| Thai patient | 316c | 17 | SRR2975745 | CAZr (64) | --- | PenA<sub>P173S</sub> | [20, 21] |
| Thai patient | 354e | 78 | AHJD00000000.1 | CAZi (6) | --- | penA -78G>A | [7] |
|  |  |  |  | SXTi (3) | BpeT structural variant<sup>‡</sup> | Ptr1<sub>R21fs</sub> |  |
| Australian patient | Bp1651 | 880 | SRR2102060 | CAZr (≥128) | penA -78G>A | PenA<sub>D245G</sub> | [28] |
|  |  |  |  | Unusual sensitivity to aminoglycosides (GEN, KAN) and macrolides (AZM) | --- | AmrB<sub>A254fs</sub> |  |
|  |  |  |  | DOXr (16) | --- | BPSL3085<sub>A88fs</sub> |  |
|  |  |  |  | AMCr (64/32) | penA -78G>A | PenA<sub>S78F</sub> |  |
| Pre-DPMS 89 | MSHR0052 | 722 | SRR5818275 | MEMr (8) | --- | AmrR<sub>E190*</sub> | [6] |
|  |  |  |  | DOXr (48) | AmrR<sub>E190*</sub> | BPSL3085<sub>V211M</sub> |  |
| Australian patient | MSHR0292 | 236 | SRR4254580 | DOXr (16) | AmrR<sub>S174P</sub> | BPSL3085<sub>V40A</sub> | [26] |
| P215 | MSHR0663 | 36 | SRR2887062 | SXTr (≥32) | BpeT<sub>H278Y</sub> | Ptr1<sub>R21fs</sub> | [6]; This study |
|  | MSHR0937 | 36 | SRR2886988 | AMCi (12/6), MEMr (6) | --- | BpeR<sub>D176A</sub> |  |
|  |  |  |  | SXTi (3) | BpeR<sub>D176A</sub> | MetF<sub>Q142*</sub> |  |
| P179 | MSHR0678 | 114 | SRR6075118 | MEMr (3) | --- | AmrR<sub>E21D</sub> | [6] |
|  | MSHR0800 | 114 | SRR6075115 | MEMr (6), DOXi (8) | --- | BpeR<sub>L85fs</sub> |  |
| P337 | MSHR1226 | 333 | SRR9598635 | CAZr (≥256) | --- | PenA<sub>C75Y</sub> | [20] |
|  | MSHR1300 | 333 | SRR6075114 | CAZr (≥256) | penA -78G>A | PenA<sub>C75Y</sub> | [20] |
|  |  |  |  | MEMr (4) | --- | AmrR<sub>K13fs</sub> | [6] |
|  |  |  |  | SXTi (3) | --- | AmrR<sub>K13fs</sub> | This study |
| **P595** | MSHR3683 | 144 | SRR11678542 | DOXi (12) | --- | BPSL3085<sub>A88fs</sub> | [9, 28] |
| **P608** | MSHR4083 | 36 | SRR2887030 | SXTr (24) | AmrR<sub>ΔA153-D156</sub> | Dut<sub>G91A</sub> | [6] |
|  |  |  |  | MEMr (6) | --- | AmrR<sub>ΔA153-D156</sub> |  |
| **CF6** | MSHR5654 | 1040 | SRR3404570 | CAZr (≥256) | PenA 30x CNV | PenA<sub>C75Y</sub> | [9] |
|  |  |  |  | CIPr (≥32) | --- | GyrA<sub>Y77S</sub> |  |
|  |  |  |  | SXTr (≥32) | BpeT<sub>T314fs</sub> | Ptr1<sub>R21fs</sub> |  |
| **CF9** | MSHR5665 | 252 | SRR3404582 | SXTr (≥32) | Ptr1<sub>W116R</sub>;<br>MetF<sub>Q28P</sub> | Dut<sub>V77A</sub> | [9, 23] |
|  |  |  |  | DOXi (6) | --- | BPSL3085<sub>A88fs</sub> |  |
|  | MSHR5666 | 252 | SRR3404597 | DOXs (1)<sup>#</sup> | --- | BPSL3085<sub>A88fs</sub> |  |
|  |  |  |  | SXTr (≥32) | Ptr1<sub>W116R</sub>;<br>MetF<sub>Q28P</sub> | Dut<sub>V77A</sub> |  |
|  | MSHR5667 | 252 | SRR3404598 | SXTr (≥32) | MetF<sub>N162T</sub>;<br>AmrR<sub>L132P</sub> | Dut<sub>N99S</sub> |  |
|  |  |  |  | MEMr (4) | --- | AmrR<sub>L132P</sub> |  |
|  |  |  |  | DOXr (48) | AmrR<sub>L132P</sub> | BPSL3085<sub>A88fs</sub> |  |
|  | MSHR5669 | 252 | SRR3404599 | DOXi (6) | --- | BPSL3085<sub>A88fs</sub> |  |
|  |  |  |  | SXTr (≥32) | Ptr1<sub>W116R</sub>;<br>MetF<sub>Q28P</sub> | Dut<sub>V77A</sub> |  |
| **P726** | MSHR6755 | 975 | SRR6075122 | MEMr (3) | --- | AmrR<sub>ΔV60-C63</sub> | [6] |
| **P797** | MSHR7587 | 437 | SRR6075129 | MEMr (4) | --- | AmrR<sub>G30D</sub> | [6] |
|  | MSHR7929 | 437 | SRR6075126 | SXTr (4) | AmrR<sub>G30D</sub> | BPSL2263<sub>ΔD110-G116</sub> |  |
|  |  |  |  | MEMr (4) | --- | AmrR<sub>G30D</sub> |  |
| **CF11** | MSHR8441 | 46 | SRR3382162 | CAZi (12) | --- | PenA 10x CNV | [6, 9] |
|  |  |  |  | SXTr (≥32) | AmrR<sub>ΔV62-H223</sub><sup>C</sup> | Ptr1<sub>A22-G23ins_R-R-A</sub> |  |
|  |  |  |  | Decreased DOX susceptibility (4) | AmrR<sub>ΔV62-H223</sub><sup>C</sup> | BPSL3085<sub>S130L</sub> |  |
|  |  |  |  | Decreased MEM susceptibility (2) | --- | AmrR<sub>ΔV62-H223</sub><sup>C</sup> |  |
|  | MSHR8442 | 46 | SRR3404603 | SXTr (≥32) | AmrR<sub>ΔV62-H223</sub><sup>C</sup> | Ptr1<sub>A22-G23ins_R-R-A</sub> |  |
|  |  |  |  | DOXi (8) | AmrR<sub>ΔV62-H223</sub><sup>C</sup> | BPSL3085<sub>S130L</sub> |  |
|  |  |  |  | MEMr (3) | --- | AmrR<sub>ΔV62-H223</sub><sup>c</sup> |  |
| **Non-DPMS**<br>**QP09** | MSHR8481 | 1378 | SRR6075123 | MEMr (6) | --- | AmrR<sub>ΔA70-H223</sub><sup>d</sup> | [6] |
| **P989** | MSHR9021<sup>a</sup> | 132 | SRR6075127 | MEMr (3) |  | AmrR<sub>S166P</sub>; AmrR<sub>A145fs</sub><sup>b</sup> | [6] |
| **Strains with unusual antibiotic sensitivity** |  |  |  |  |  |  |  |
| **P314** | MSHR1043<sup>§</sup> | 131 | SRR6380769 | Unusual GEN sensitivity | --- | AmrA<sub>L247fs</sub> | [14] |
| **Malaysian patient** | MSHR5089<sup>§</sup> | 881 | SRR2975737 | Unusual sensitivity to aminoglycosides (GEN, KAN) and macrolides (AZM) | --- | AmrB<sub>T368R</sub> | [17] |
| **Strains with stepwise variants** |  |  |  |  |  |  |  |
| **P179** | MSHR0535 | 114 | SRR6075120 | DOXs (4)<sup>§</sup> | BPSL3085<sub>R104fs</sub> | --- | [6] |
| **Thai patient** | 354e | 78 | AHJD00000000.1 | AMCs (4/2) | penA -78G>A | --- | [7] |

Abbreviations: AZM, azithromycin; CAZ, ceftazidime; CIP, ciprofloxacin; CNV, copy-number variation; DOX, doxycycline; GEN, gentamicin; KAN, kanamycin; MEM, meropenem; MIC, minimum inhibitory concentration; SXT, co-trimoxazole
<sup>‡</sup>i, intermediate; r, resistant; s, sensitive
<sup>†</sup>Where applicable
<sup>‡</sup>Encodes an 800kb inversion that direct affects *llpE*, the intervening gene between the BpeT transcriptional regulator and its BpeEF-OprC efflux pump [7]
<sup>#</sup>False positive; isolate remained sensitive despite encoding a known AMR determinant. Cause of reversion unknown
<sup>§</sup>Strains MSHR1043 [14] and MSHR5089 [17] encode unusual antimicrobial susceptibility, and are susceptible to all five clinically-relevant antibiotics (i.e. AMC, CAZ, DOX, MEM, SXT), and strains MSHR0535 and 354e encode stepwise variants that elevate the DOX and AMC MICs, respectively, but do not exceed the established resistance cut-off for these antibiotics
<sup>a</sup>MSHR9021 was intentionally sequenced from a potentially mixed population to capture population diversity. Two AmrR mutants, AmrR<sub>S166P</sub> and AmrR<sub>A145fs</sub>, were present at ratios of ~66% and 33%, respectively [6]. In the non-mixture mode, only the dominant variant, AmrR<sub>S166P</sub>, is detected.
<sup>b</sup>Frameshift indel shortens protein length from 223 to 183 residues
<sup>c</sup>Frameshift indel increases protein length from 223 to 285 residues
<sup>d</sup>Frameshift indel shortens protein length from 223 to 117 residues

**Table 2.** *Burkholderia pseudomallei* strains phenotypically confirmed to be sensitive towards the five clinically-relevant antibiotics, and associated genome data.

| Patient ID | Isolate | ST | Genome accession | Antibiotic MIC (µg/mL)* |  |  |  |  | Reference/s |
| --- | --- | --- | --- | --- | --- | --- | --- | --- | --- |
|  |  |  |  | AMC | CAZ | DOX | MEM | SXT |  |
| Australian patient | MSHR0293 | 236 | SRR4254579 | 2/1 | 1 | 1 | 0.5 | 0.4 | [26] |
| P179 | MSHR0492 | 114 | SRR6075119 | 1.5/0.75 | 1.5 | 1 | 1.5 | 1 | [6] |
|  | MSHR0934 | 114 | SRR6075116 | 2/1 | 2 | 1 | 0.75 | 1 |  |
| P337 | MSHR1141 | 333 | SRR2975732 | 1.5/0.75 | 1.5 | 1 | 0.75 | 0.75 | [20] |
| P608 | MSHR3763 | 36 | SRR2887021 | 4/2 | 2 | 0.75 | 0.75 | 3 | [6] |
| P726 | MSHR5864 | 975 | SRR6075121 | 3/1.5 | 1.5 | 1 | 0.75 | 1.5 | [6] |
| P797 | MSHR6522 | 437 | SRR6075128 | 2/1 | 1.5 | 1 | 0.5 | 1.5 | [6] |
| CF1 | MSHR0913 | 279 | SRR3404575 | 2/1 | 3 | 1 | 1 | 0.5 | [9] |
|  | MSHR1053 | 279 | SRR3404578 | 1.5/0.75 | 2 | 1 | 0.5 | 1 |  |
| Malaysian patient | MSHR5093 | 881 | SRR2975738 | 6/3 | 4 | 1.5 | 1 | 3 | [17] |
| Malaysian patient | MSHR5104 | 881 | SRR2975740 | 2/1 | 0.75 | 1 | 0.19 | 0.75 | [17] |
| CF6 | MSHR5651 | 1040 | SRR3381886 | 1.5/0.75 | 1.5 | 0.38 | 0.5 | 0.75 | [9, 23] |
| CF9 | MSHR5662 | 252 | SRR3381885 | 2 | 2 | 2 | 0.75 | 0.5 | [9, 23] |
|  | MSHR5670<sup>#</sup> | 252 | SRR3404600 | 1.5/0.75 | 2 | 2 | 0.75 | 0.5 |  |
| CF10 | MSHR8438 | 442 | SRR3382015 | 2/1 | 3 | 1 | 1 | 0.25 | [9, 23] |
|  | MSHR8440 | 442 | SRR3404601 | 2/1 | 3 | 1 | 0.75 | 0.38 |  |

Abbreviations: AMC, amoxicillin-clavulanate; CAZ, ceftazidime; DOX, doxycycline; MEM, meropenem;
ST, multilocus sequence type; SXT, co-trimoxazole
\*According to Etesting

**Table 3.** ARDaP prediction in three meropenem-resistant *Burkholderia pseudomallei* isolates with previously unknown antimicrobial resistance (AMR) determinants.

| Strain ID | ARDaP-predicted AMR determinant | MEM MIC (µg/mL) | Reference/s |
| --- | --- | --- | --- |
| MSHR1058 | AmrR<sub>ΔP81-H223</sub><sup>a</sup> | 12 (MEM) | [6]; This study |
| MSHR1174 | AmrR<sub>G149fs</sub><sup>b</sup> | 6 (MEM) | This study |
| MSHR8777 | AmrR<sub>ΔA128-H223</sub><sup>c</sup> | 4 (MEM) | This study |

<sup>a</sup>Previously undescribed 3' *amrR* deletion shortens protein length from 223 to 130 residues.
<sup>b</sup>Previously undescribed 11bp insertion shortens protein length from 223 to 178 residues.
<sup>c</sup>Previously undescribed 3' *amrR* insertion increases protein length from 223 to 621 residues.

### Culturing, WGS, and genome assembly

*B. pseudomallei* culture, DNA extraction, and WGS were performed as described elsewhere [9]. Genomic data for MSHR1058, MSHR1174, and MSHR8777 were uploaded to the NCBI Sequence Read Archive database (BioProject PRJNA641249). Accession numbers for all other genomic data are listed in Tables 1 and 2. For genomes lacking a publicly-available assembly, MGAP v1.1 (https://github.com/dsarov/MGAP--Microbial-Genome-Assembler-Pipeline) was used, with archetypal strain K96243 (RefSeq accessions NC_006350.1 and NC_006351.1) provided as the scaffolding reference.

### Minimum Inhibitory Concentrations (MICs)

MICs were determined using Etests (bioMérieux, Murarrie, Australia). Sensitive, intermediate and resistant cut-offs were based on the Clinical and Laboratory Standards Institute (CLSI) M100-S17 guidelines for *B. pseudomallei* (≤8/4, 16/8, and ≥32/16 μg/mL for AMC; ≤8, 16, ≥32 μg/mL for CAZ; ≤4, 8, ≥16 μg/mL for DOX and IPM; ≤2/38, nil, ≥4/76 μg/mL for SXT). CLSI guidelines do not list MEM for *B. pseudomallei*; however, based on prior work [15, 16], and recent proposed EUCAST breakpoints for *B. pseudomallei*, we categorised MEMr as ≥3 μg/mL. Likewise, the CLSI guidelines do not list gentamicin (GEN) MIC values for *B. pseudomallei* due to almost ubiquitous resistance (>16 μg/mL) towards this antibiotic; however, there are notable exceptions [14, 17]. We chose a GEN-sensitive cut-off of ≤4 μg/mL, which also reflects those strains unable to grow on Ashdown’s agar, a selective medium for *B. pseudomallei* isolation that contains 4 μg/mL GEN.

### AMR software parameters

The default RGI v5.1.0 database parameters of CARD v3.0.9 (https://card.mcmaster.ca/analyze/rgi; accessed 25Jun20), and ARIBA v2.14.5 (https://github.com/sanger-pathogens/ariba), were examined for performance across the *B. pseudomallei* genomes.

### ARDaP AMR database construction

ARDaP is available at: https://github.com/dsarov/ARDaP. All reported *B. pseudomallei* AMR determinants, including stepwise AMR mutations and unusual antimicrobial susceptibility mutations (Table 1; Table S1), were annotated relative to K96243. The AMR determinants (as of version 1.7) are summarised in an SQLite database (Table S1; most up-to-date version available at: https://github.com/dsarov/ARDaP/tree/master/Databases/Burkholderia_pseudomallei_k96243). Briefly, CAZ resistance (CAZr) is caused by altered PenA β-lactamase substrate specificity [18–21], *penA* upregulation [7, 20, 22, 23] (including CNVs [9]), or loss of penicillin-binding protein 3 [24]; AMC resistance (AMCr) is caused by *penA* upregulation [14, 20]; MEMr is caused by AmrAB-OprA, BpeAB-OprB, or BpeEF-OprC resistance-nodulation-division (RND) multidrug efflux pump regulator loss-of-function [6]; SXT resistance (SXTr) is caused by cumulative mutations in core metabolism pathways coupled with AmrAB-OprA, BpeAB-OprB, or BpeEF-OprC RND efflux pump regulator loss-of-function [6, 9, 25]; and DOX resistance (DOXr) is caused by loss-of-function mutations within the SAM-dependent methyltransferase gene, *BPSL3085*, often in combination with AmrAB-OprA, BpeAB-OprB, or BpeEF-OprC regulator loss-of-function [26]. Our *B. pseudomallei* ARDaP database also includes AmrA and AmrB mutants that are associated with unusual aminoglycoside and macrolide susceptibility [14, 17]. To avoid poor-quality WGS data or incorrect species assignments, the database also includes two conserved genetic targets (Table S1) found only in this bacterium; strains lacking these loci are flagged for further user assessment.

### ARDaP algorithm

To achieve high-quality variant calls, ARDaP incorporates several tools into its workflow (full list available at: https://github.com/dsarov/ARDaP). In addition to an organism-specific SQLite database (https://github.com/dsarov/ARDaP/tree/master/Databases), ARDaP requires WGS data, either genomes or metagenomes in paired-end Illumina v1.8+ FASTQ format, or assembled genomes in FASTA format, as input (Figure 1). For genomes in FASTA format, ARDaP first converts to synthetic Illumina v1.8+ reads. For genomes in FASTQ format, ARDaP performs quality filtering followed by optional random down-sampling to a user-defined coverage (default=50x) to permit more rapid analysis. ARDaP then performs comparative genomic analysis to identify AMR determinants by mapping reads against an annotated reference, followed by alignment processing, SNP and indel identification, coverage assessment, and structural variant identification. High-quality genetic variants (SNPs, indels [<50bp], CNVs, gene gain, and gene loss or truncation) are then annotated. ARDaP next interrogates two databases: i) a customisable CARD [5] database is screened to identify horizontally-acquired AMR genes and to ignore conserved genes that do not confer AMR, and ii) a bespoke AMR determinant database (in this study, a *B. pseudomallei* database) containing species specific AMR determinants. ARDaP databases are created in SQLite and can be readily updated as additional AMR determinants are identified. This database also accommodates stepwise mutations and AMR conferred by ≥2 mutations. Finally, ARDaP can predict AMR by identifying novel high-consequence mutations (i.e. those resulting in a frameshift or nonsense mutations) in known AMR genes. These putative mutants can then be flagged for further investigation with phenotypic AMR testing. ARDaP outputs are presented in a comprehensive, human-readable report (Figure 3), which has been modelled from the report structure described by Crisan and colleagues.

**Figure 1.**
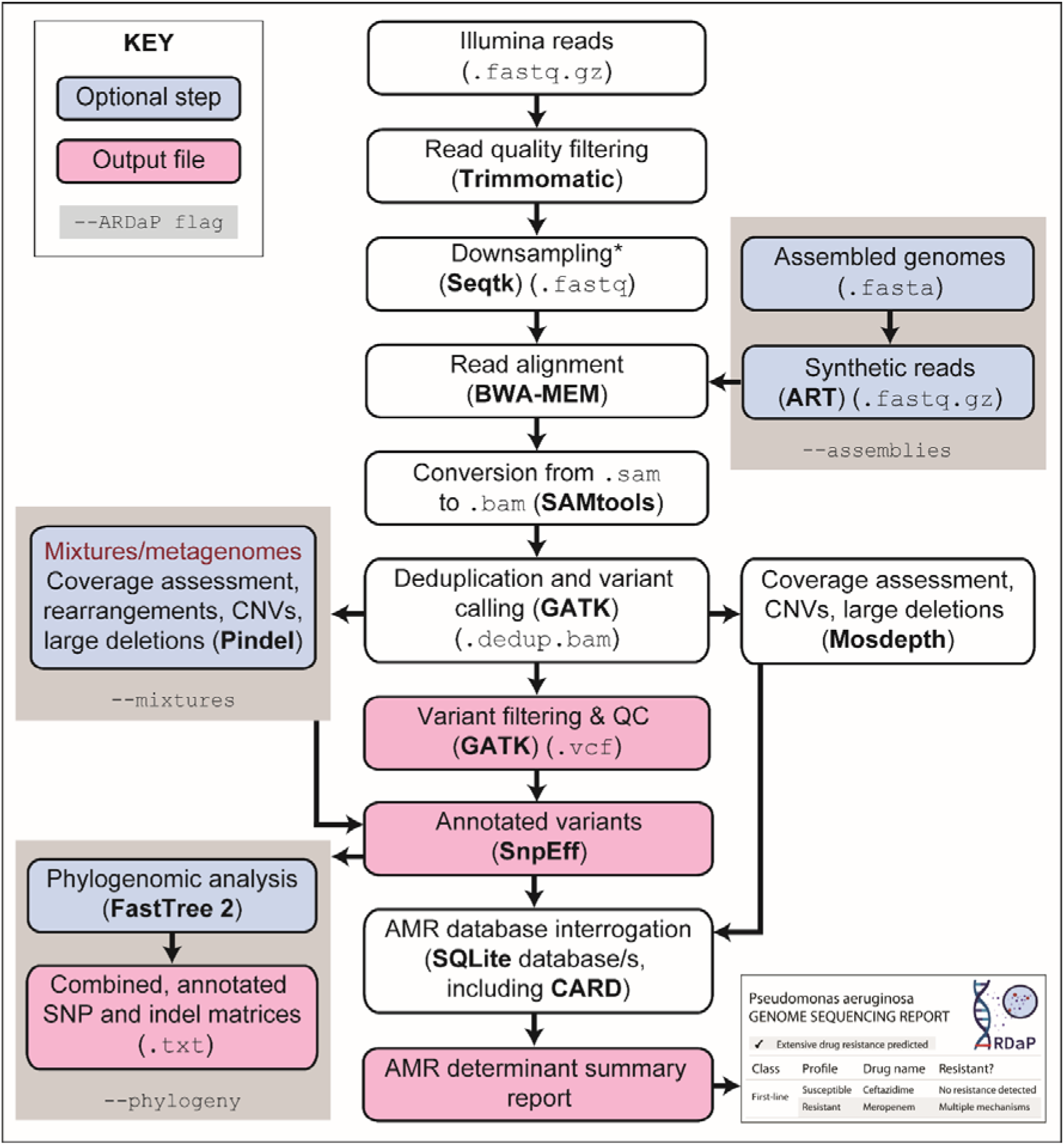
ARDaP pipeline. The user inputs assembled genome/s or raw sequencing reads, and a reference genome sequence. ARDaP then performs read alignment, read processing, deduplication, and variant identification. An optional phylogenetic analysis is also performed (if specified). Coverage assessment is undertaken on either single or mixed genomes (if specified); genetic variants are then annotated and antimicrobial resistance database/s interrogated. Finally, ARDaP produces a summary report of antimicrobial resistance determinants for each strain (Figure 2). *Downsampling is carried out by default but can be turned off using the `--size 0` flag in ARDaP.

### Mixture detection

ARDaP incorporates a minor allelic variant analysis function to permit variant identification from mixed genomes/metagenomes, enabling the detection of emerging AMR determinants (down to 5% abundance). Minor-variant SNPs and indels are identified using the ploidy-aware HaplotypeCaller tool in GATK v4.1; deletions, CNVs, and structural mutations are identified with the ploidy-aware function of Pindel. *B. pseudomallei* strains with known AMR status were mixed at ratios of 5% increments ranging from 5:95 to 95:5, to 55-60x total depth. Two mixtures were created: MSHR0913 (sensitive to all clinically-relevant antibiotics) and MSHR0937 (MEMr; intermediate resistance to AMC and SXT), and MSHR0913 and MSHR8441 (SXTr; decreased susceptibility to CAZ, MEM, and DOX). MSHR0937 and MSHR8441 were chosen as they represent a wide spectrum of clinically-relevant AMR and mutation types (Table 4).

**Table 4.**
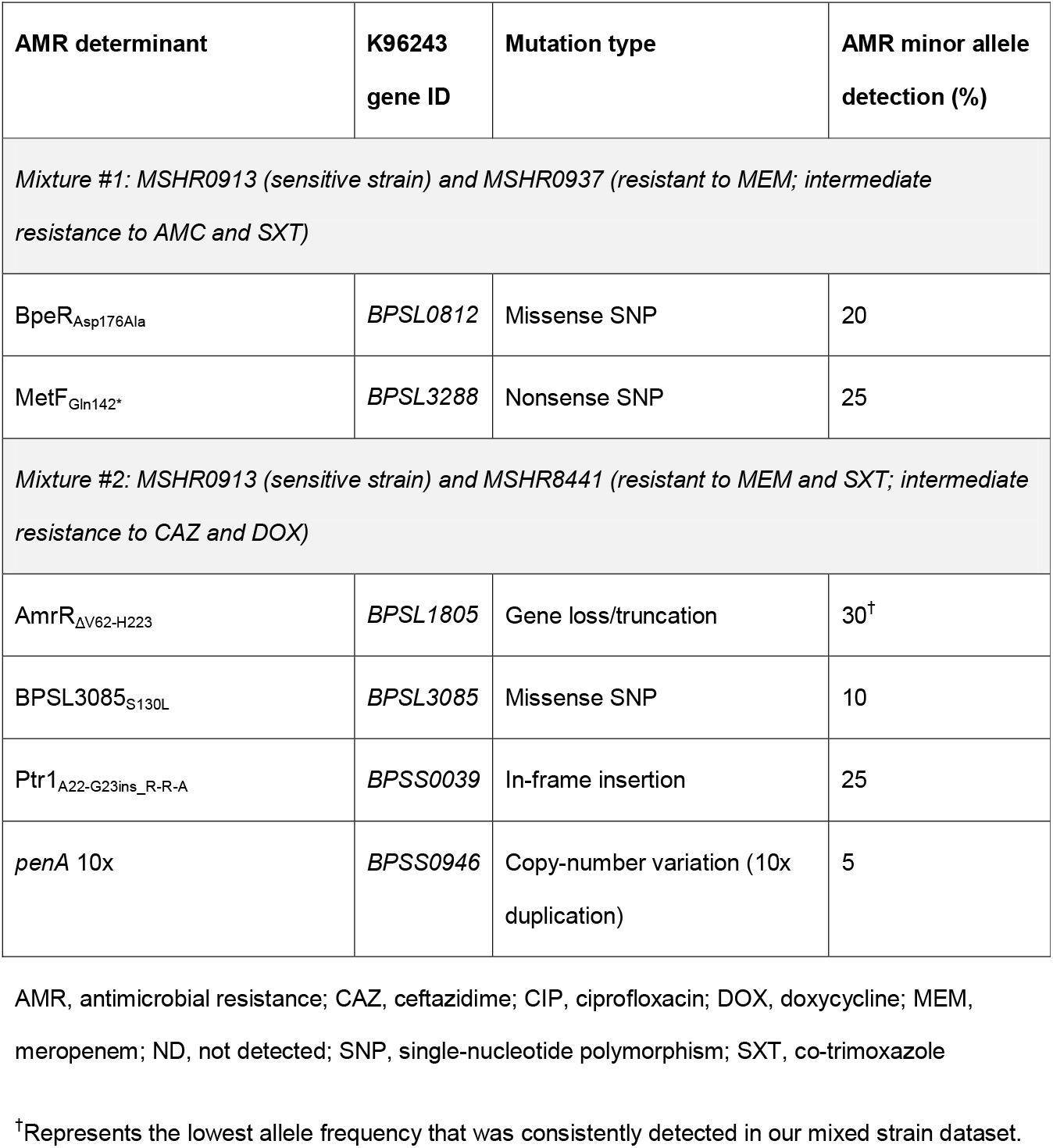
ARDaP detection limits for AMR determinants in synthetic mixtures.

| AMR determinant | K96243<br>gene ID | Mutation type | AMR minor allele<br>detection (%) |
| --- | --- | --- | --- |
| *Mixture #1: MSHR0913 (sensitive strain) and MSHR0937 (resistant to MEM; intermediate resistance to AMC and SXT)* |  |  |  |
| BpeR<sub>Asp176Ala</sub> | BPSL0812 | Missense SNP | 20 |
| MetF<sub>Gln142*</sub> | BPSL3288 | Nonsense SNP | 25 |
| *Mixture #2: MSHR0913 (sensitive strain) and MSHR8441 (resistant to MEM and SXT; intermediate resistance to CAZ and DOX)* |  |  |  |
| AmrR<sub>ΔV62-H223</sub> | BPSL1805 | Gene loss/truncation | 30<sup>†</sup> |
| BPSL3085<sub>S130L</sub> | BPSL3085 | Missense SNP | 10 |
| Ptr1<sub>A22-G23ins_R-R-A</sub> | BPSS0039 | In-frame insertion | 25 |
| *penA* 10x | BPSS0946 | Copy-number variation (10x duplication) | 5 |

AMR, antimicrobial resistance; CAZ, ceftazidime; CIP, ciprofloxacin; DOX, doxycycline; MEM, meropenem; ND, not detected; SNP, single-nucleotide polymorphism; SXT, co-trimoxazole
<sup>†</sup>Represents the lowest allele frequency that was consistently detected in our mixed strain dataset.

## Results and Discussion

### Performance of existing AMR tools in *B. pseudomallei*

The validated dataset of 47 *B. pseudomallei* isolates was used to assess the performance and capacity of existing AMR tools to identify AMR determinants in AMR but not antimicrobial-sensitive strains. According to CARD, all 47 genomes were found to harbour AMR determinants; however, all determinants corresponded with conserved genes in *B. pseudomallei* (Table S2). In addition, CARD failed to identify any clinically relevant AMR determinants in the 25 AMR strains. This finding is consistent with *B. pseudomallei* acquiring AMR through chromosomal mutation rather than gene gain, and an inability of current CARD software to identify AMR variants conferred by indels, CNVs, or gene loss/truncation. ARIBA outperformed CARD due to its ability to include missense (although not nonsense) SNPs in its database construction and to identify SNPs and indels in its report outputs. However, we encountered several shortcomings with ARIBA; notably, an inability to differentiate AMR determinants from natural variation, cumbersome and labour-intensive input file requirements, restrictions on database construction (e.g. reference genomes with indels and nonsense mutations cannot be included), and an inability to identify CNVs. In our hands, ARIBA provided comparable information to variant reporting outputs generated by comparative genomic pipelines. However, such efforts require extensive domain-specific knowledge to accurately uncover and interpret AMR determinants. In contrast, ARDaP uses a single and standardised SnpEff-based input, can differentiate natural gene variation from known and putative AMR determinants, can detect CNVs, and provides a user-friendly output that does not require domain-specific knowledge to interpret outputs.

### ARDaP development and performance in *B. pseudomallei*

Due to shortcomings in existing AMR software, ARDaP was designed to both identify known AMR determinants and to ignore non-causal genetic variants (Table 2). ARDaP correctly identified all *B. pseudomallei* AMR determinants (Table 1); however, when tested against the 47 validated isolates, one false negative and four false positives were identified. The false negative result (SXT sensitivity rather than decreased SXT susceptibility) was due to an 800kb inversion in 354e that directly impacts *llpE*, a gene that resides between *bpeT* and *bpeEF-oprC* [7]. BpeT is a LysR-type transcriptional regulator that controls expression of BpeEF-OprC [27]. This inversion was not detected due to limitations in short-read Illumina data that led to this structural variant not easily being identified by Pindel, coupled with the occurrence of this inversion outside of an AMR determinant.

The first of the false positives, chronic cystic fibrosis (CF) isolate MSHR5654, was predicted to be MEMr due to the presence of BpeT_Thr314fs_. However, Etesting showed MEM sensitivity (2 μg/mL) in this strain, just below the MEMr threshold (≥3 μg/mL) [9, 23]. Although alterations in BpeT have been putatively linked with MEMr in MSHR1300 (4 μg/mL) [6] and 354e (6 μg/mL) [7], the role of BpeT mutations in conferring MEMr is contentious [25]. In support of this notion, MSHR1300 also encodes AmrR_K13fs_, a TetR-family *cis*-acting repressor of the AmrAB-OprA RND efflux pump, which likely causes MEMr in its own right [6], and in 354e, the ~800kb inversion likely also affects other AMR-conferring genes besides *bpeT*. As such, the *B. pseudomallei* ARDaP database was updated to flag *bpeT* variants as stepwise mutations rather than solely conferring MEMr (Table 1; Table 2), thereby correcting the original false-positive call for MSHR5654. This issue highlights the complexity of unravelling AMR determinants, and in this case, the need for additional work to determine a role, if any, for *bpeT* mutations in conferring MEMr.

The remaining false-positive strains, MSHR5666, MSHR5669, and MSHR5670, were all obtained from a chronically-infected CF patient, CF9 [9]. All encode a SAM-dependent methyltransferase truncation (BPSL3085_A88fs_) [23] and were predicted by ARDaP to be DOXr. We also observed BPSL3085_A88fs_ in an unrelated DOXr chronic CF strain, Bp1651 [28] (Table 1). BPSL3085 mutations confer DOXr by altering ribosomal methylation patterns [9, 26]. However, all three strains remained DOX-sensitive (1.5 μg/mL) despite other CF9 strains encoding BPSL3085_A88fs_ and being DOXr (MSHR5665: MIC=6 μg/mL; MSHR5667: MIC=48 μg/mL; Table 1) [9]. The higher DOX MIC in MSHR5667 is attributable to a second mutation (AmrR_L132P_; Table 1). We postulate that MSHR5666, MSHR5669, and MSHR5670 encode an unidentified mutation that reverts them to a DOX-sensitive phenotype. Notably, all longitudinal CF9 isolates, including MSHR5666 and MSHR5669, encode *mutS* mutations, resulting in a hypermutator phenotype [9, 23]. Therefore, identifying the causal basis for this reversion is non-trivial due to the large number of mutations (range: 112-157) accrued by these hypermutator strains [9]. In addition, MSHR5670 was predicted to be SXTr due to Ptr1_W116R_, MetF_Q28P_, and Dut_V77A_ variants, yet exhibited SXT sensitivity (Table 2). The cause of SXTr reversion in this strain is also currently unknown.

### Importance of including natural genetic variation in AMR databases

Accurate prediction of novel AMR determinants requires thorough cataloguing of both confirmed AMR-causing mutations and natural variation in AMR-encoding genes to avoid false positives. As an illustration, a PenA β-lactamase missense mutation (K96243 numbering: PenA_S78F_; encoded by *BPSS0946*) has been linked to AMCr [18, 21, 28]. However, we found that PenA_S78F_ alone is unlikely to cause AMCr due to its presence in genetically diverse AMC-sensitive strains (MIC=3-4 μg/mL in strains MSHR0291, MSHR0668, MSHR0848, MSHR0911, MSHR1711, MSHR2212, MSHR3902, MSHR4797, MSHR8392, and MSHR9887). Instead, AMCr is likely conferred by both PenA_S78F_ and *penA* upregulation, the latter of which can be caused by mutations within the 5’ untranslated region [20] or *penA* CNVs [9]. We therefore included PenA_S78F_ as a putative stepwise mutation (Table 2), with an additional *penA* upregulation mutation required to confer the AMCr phenotype. In another example, we observed that both AMR and antimicrobial-sensitive strains can possess 3’-truncated *amrR* (Table 2). Multiple frameshift mutations and deletions in *amrR* are associated with MEMr [6] due to loss-of-repressor function (Figure 3). However, 3’ region mutations (residues ~210-223) do not cause MEMr (Table 2). To accommodate this natural genetic variation, we coded ARDaP to ignore these non-causal 3’ variants, thereby greatly reducing false-positive MEMr rates.

### No evidence of IPM-resistant *B. pseudomallei*

In most melioidosis treatment guidelines, IPM has been replaced by MEM due to neurotoxicity concerns.[29] However, the recent discovery of MEMr *B. pseudomallei* has resurrected IPM as a treatment option due to a lack of cross-resistance between these carbapenems [6] and exceedingly low rates of reported IPM resistance (IPMr) [30]. The one study reporting an IPMr (MIC=8 μg/mL) *B. pseudomallei* strain, Bp1651, attributed this phenotype to a PenA_T147A_ mutation (K96243 numbering: PenA_T153A_) combined with upregulation due to a promoter mutation [28]. We subsequently refuted the role of the PenA_T147A_ variant alone in conferring IPMr by identifying three genetically unrelated PenA_T153A_-encoding strains that were IPM-sensitive [6]. Further, this variant is dominant (>50%) in publicly available *B. pseudomallei* genomes, most of which exhibit wild-type antimicrobial sensitivity. Given that PenA_T147A_ occurs at a very high rate in the wild-type *B. pseudomallei* population, and none have been shown to exhibit IPMr, this mutant has not been included in our ARDaP database. However, this variant can readily be added as a stepwise AMR determinant should further evidence come to light about its role in conferring AMR.

### Reversions and unusual antimicrobial susceptibility

Aminoglycoside- and macrolide-class antibiotics are typically not included in melioidosis treatment regimens due to near-ubiquitous intrinsic resistance; indeed, GEN resistance is commonly used for *B. pseudomallei* selection [17]. However, rare cases of sensitivity have been documented, such as in ST-881 and ST-997 strains from Sarawak, Malaysian Borneo, which naturally encode AmrB_T368R_, resulting in AmrAB-OprA loss-of-function and unusual aminoglycoside and macrolide susceptibility [17]. An AmrAB-OprA loss-of-function variant has also been described in Bp1651 (AmrB_A254fs_) [28]. Although ARDaP detected *amrR* loss in MSHR1043, the co-presence of *amrA* loss (AmrA_L247fs_) resulted in reversion of MEMr to a wild-type MIC (0.75 μg/mL). This reversion also causes unusual gentamicin (MIC=1 μg/mL) [14] and presumably kanamycin and azithromycin sensitivity. Given their confounding potential, we incorporated these reversions into ARDaP to more accurately reflect the true strain phenotype. In addition, strains encoding AmrAB-OprA loss-of-function variants are at far lower risk of developing MEMr than wild-type strains, supporting longer-term MEM use in such cases. These findings highlight the value of including sensitivity-conferring variants in AMR prediction databases by increasing the antibiotic arsenal in naturally multidrug-resistant pathogens where treatment options are limited.

### AMR predictive capacity of ARDaP

We tested ARDaP’s predictive capacity to identify the causative mutation/s in three clinical MEMr strains (MSHR1058, MSHR1174, and MSHR8777; MEM MIC range: 4-12 μg/mL; Table 3) with no previously reported AMR determinants. ARDaP identified novel *amrR* mutations in each strain, all of which resulted in AmrR loss-of-function (AmrR_ΔP81-H223_ in MSHR1058; AmrR_G149fs_ in MSHR1174; AmrR_ΔA128-H223_ in MSHR8777; Figure 3). All patients received MEM treatment prior to isolate retrieval, consistent with selection pressure driving AMR. These findings provide further confirmation of the link between MEM treatment and potential treatment failure due to AmrR mutability [6], and demonstrate the value of ARDaP for predicting novel AMR determinants.

### ARDaP performance on mixed sequence data

The ARDaP algorithm is mixture-aware, an important feature for detecting emerging AMR determinants in mixed strain data (e.g. non-purified colonies, culture sweeps, clinical specimens). To assess the performance of the mixture function in ARDaP, Illumina reads from antimicrobial-sensitive and AMR *B. pseudomallei* strains (Table 4) were mixed at 5% incremental ratios ranging from 5:95 to 95:5. ARDaP identified one AMR determinant down to the lowest tested ratio of 5% minor allele frequency: a *penA* 10x CNV from MSHR8441 (Table 4). The other determinants were identified by ARDaP when present at minor allele frequencies of 10% (BPSL3085_S130L_), 20% (BpeR_Asp176Ala_). 25% (Ptr1_A22_G23ins_R-R-A_ and MetF_Gln142*_) and 30% (AmrR_ΔV62-H223_) (Table 4). Although CNVs were readily identified at low allelic frequencies, and the detection of SNPs and indels ranged from 10 to 25%, gene truncations had the lowest sensitivity due to the challenge of discriminating gene loss from Illumina depth coverage variation, coupled with inherent limitations in short-read data mapping. Next, ARDaP was tested on a previously detected AmrR mixture from strain MSHR9021, which encodes AmrR_S166P_ and AmrR_A145fs_ variants at ~66% and ~33% allele frequencies, respectively ^6^ (Table 1). ARDaP detected AmrR_S166P_ and AmrR_A145fs_ at allele frequencies of 63% and 31%, respectively, thus closely reflecting their known proportions. Overall, ARDaP confidently identified all mixtures, albeit with varying sensitivities. Further validation on specific variant mixtures is recommended when new mixtures are identified to determine their sensitivity. Deeper sequencing (e.g. 100-500x) should enable more robust mixture detection at lower allele frequencies.

### ARDaP reports

ARDaP generates an easy-to-interpret report that summarises the AMR determinants and associated antibiotic phenotype/s for each genome. This report summarises AMR findings for first-line, second-line, and tertiary antibiotics (Figure 2), along with instances of unusual antibiotic susceptibility, and has been designed to prioritise a clinical workflow. This report also lists stepwise AMR determinants, thereby informing early treatment shifts that mitigate the risk of AMR emergence and fixation. This easy-to-interpret report represents a major improvement over current software for AMR annotation such as ARIBA and CARD, both of which require an intimate understanding of AMR determinants to correctly interpret outputs and to ignore natural genetic variation. The AMR report produced by ARDaP represents an important step towards the incorporation of WGS as a routine tool for guiding best-practice AMR stewardship and personalised treatment regimens in the clinical diagnostic setting.

**Figure 2.**
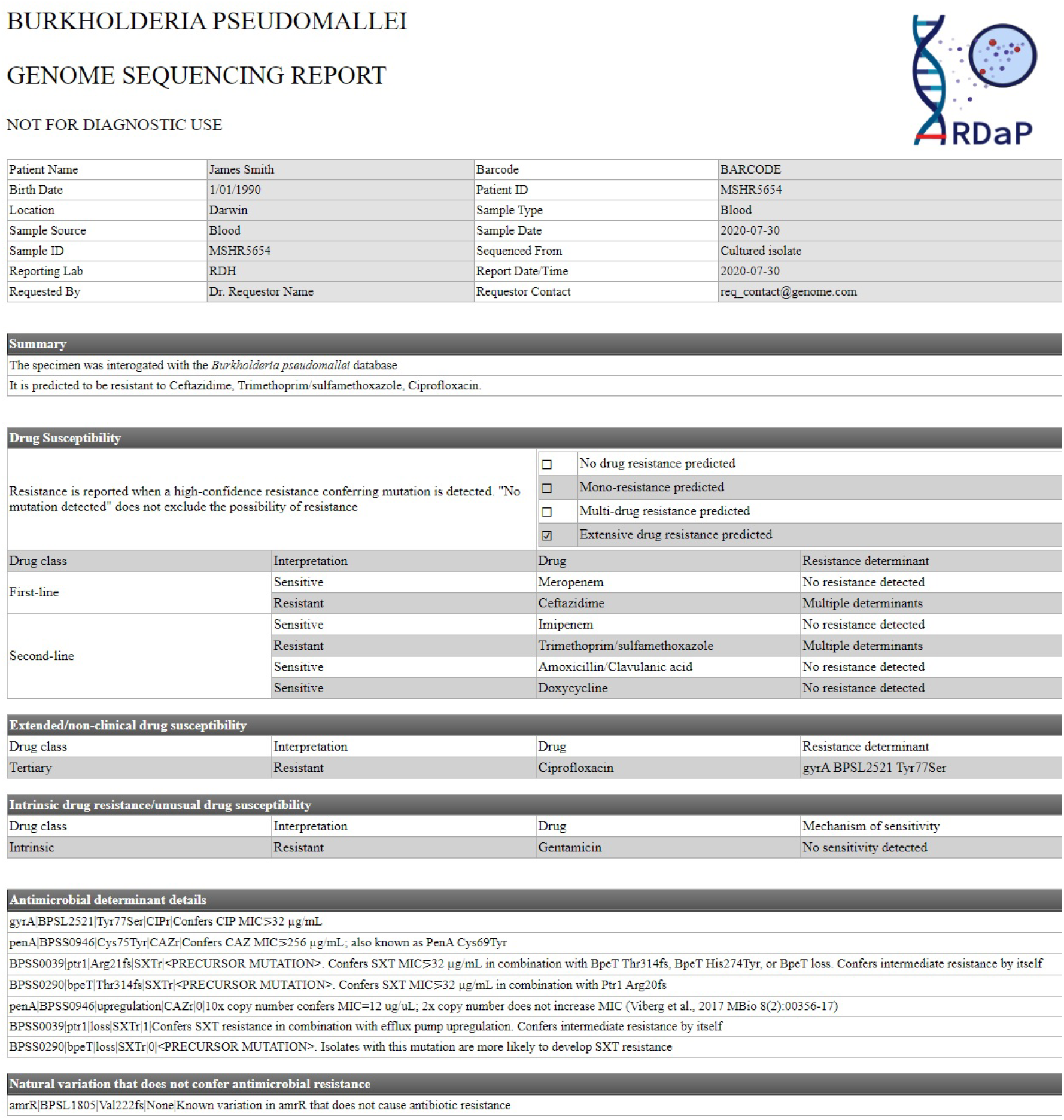
Example antimicrobial resistance (AMR) summary report produced by ARDaP. The final step in the ARDaP pipeline is the production of a user-friendly report that summarises patient and sample details, confirms that the given isolate is the expected species (in this case, *Burkholderia pseudomallei*), identifies any predicted AMR (including what mutation/s has/have been detected) and what antibiotic/s have been affected, identifies unusual antimicrobial sensitivity and natural variation that does not confer AMR, and identifies stepwise mutations that lower the barrier to AMR development.

**Figure 3.**
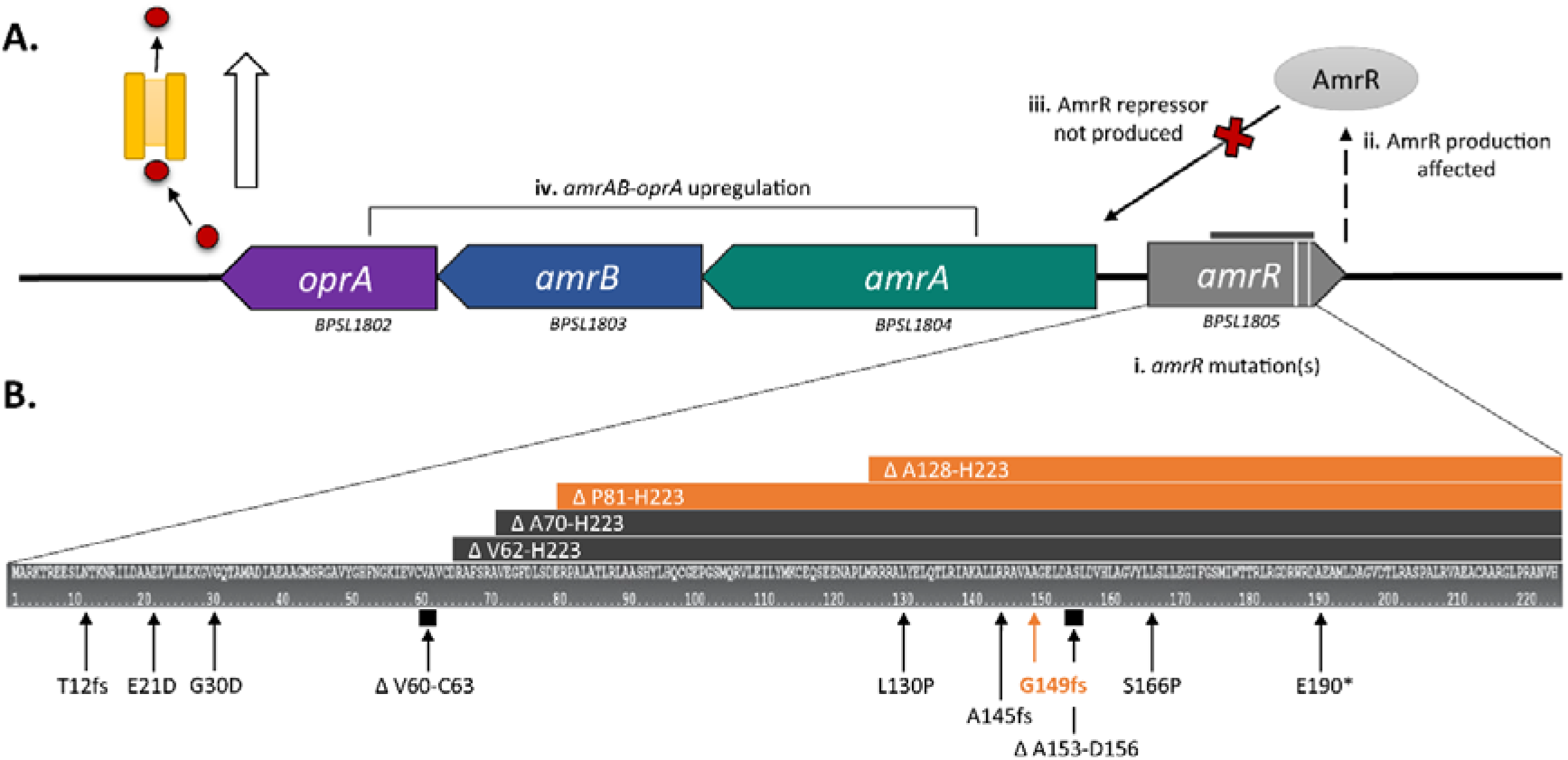
Operon organisation of the *Burkholderia pseudomallei* AmrAB-oprA resistance-nodulation-division efflux pump and loss-of-function mutations in its TetR-type regulator, AmrR. A. Transcriptional organisation of the *amrR* (*BPSL1805*), *amrA* (*BPSL1804*), *amrB* (*BPSL1803*) and *oprA* (*BPSL1802*) operon, and summary of how (i) *amrR* mutations cause (ii) loss-of-function of AmrR, which (iii) no longer represses expression of the resistance-nodulation-division AmrAB-OprA efflux pump, resulting in (iv) efflux pump over-expression and resistance to meropenem and aminoglycoside antibiotics. B. Distribution and annotation of *amrR* mutations. Eleven previously observed *amrR* mutations (in black) [6] have been augmented with three novel mutations identified in the current study (orange); AmrR_G149fs_, AmrR_ΔP81-H223_, and AmrR_ΔA128-H223_, all of which cause *amrR* loss-of-function, resulting in efflux pump overexpression and meropenem resistance.

## Acknowledgements

We thank Associate Professor Rob Baird and the microbiology staff at Royal Darwin Hospital for their support and expertise in identifying and characterising *B. pseudomallei* isolates, and Vanessa Rigas, Glenda Harrington, and Mark Mayo for isolate inventory support.

## Author contributions

DSS conceived of the study; EPP and DSS designed the study; JRW, EPP, and DSS generated laboratory data and performed laboratory analyses; EJS and DSS wrote the software; DEM, EPP, and DSS performed data analysis, literature searches, figure generation, software testing, and feature development; BJC provided isolates; DEM, EPP, and DSS wrote the manuscript; and BJC, EPP, and DSS obtained funding for the study. All authors approved of the final manuscript.

